# Can we sort states of environmental DNA (eDNA) from a single sample?

**DOI:** 10.1101/2023.01.13.523867

**Authors:** Anish Kirtane, Hannah Kleyer, Kristy Deiner

## Abstract

Environmental DNA (eDNA) once shed can exist in numerous states with varying behaviors including degradation rates and transport potential. In this study we consider three states of eDNA: 1) a membrane-bound state referring to DNA enveloped in a cellular or organellar membrane, 2) a dissolved state defined as the extracellular DNA molecule in the environment without any interaction with other particles, and 3) an adsorbed state defined as extracellular DNA adsorbed to a particle surface in the environment. Capturing, isolating, and analyzing a target state of eDNA provides utility for better interpretation of eDNA degradation rates and transport potential. While methods for separating different states of DNA have been developed, they remain poorly evaluated due to the lack of state-controlled experimentation. We evaluated the methods for separating states of eDNA from a single sample by spiking DNA from three different species to represent the three states of eDNA as state-specific controls. We used chicken DNA to represent the dissolved state, cultured mouse cells for the membrane-bound state, and salmon DNA adsorbed to clay particles as the adsorbed state. We performed the separation in three water matrices, two environmental and one synthetic, spiked with the three eDNA states. The membrane-bound state was the only state that was isolated with minimal contamination from non-target states. The membrane-bound state also had the highest recovery (54.11 ± 19.24 %), followed by the adsorbed state (5.08 ± 2.28 %), and the dissolved state had the lowest total recovery (2.21 ± 2.36 %). This study highlights the potential to sort the states of eDNA from a single sample and independently analyze them for more informed biodiversity assessments. However, further method development is needed to improve recovery and reduce cross-contamination.

## 1. Introduction

Environmental DNA (eDNA) is DNA that can be extracted from environmental samples such as water, soil, or air without first isolating any target organisms (Taberlet et al., 2012). Environmental DNA is expected to be a complex mixture of DNA from many organisms and potentially reside in different states due to the many sources from which eDNA can arise (e.g., mucus, tissues, whole single-celled organisms, etc; (Mauvisseau et al., 2022; Rodriguez-Ezpeleta et al., 2021). Environmental DNA encapsulated within a cell or organelle (e.g., nucleus or mitochondria) is considered to be membrane-bound DNA (Mauvisseau et al., 2022; Nagler et al., 2022). After membrane lysis, the DNA becomes extracellular and can be further categorized into two states when in water; dissolved DNA having no interactions with other particles and adsorbed DNA referring to DNA chemically or physically bound to particles (Mauvisseau et al., 2022; Nagler et al., 2022). Thus, at a minimum, eDNA from environmental samples is likely to exist in at least three states (i.e., membrane-bound, dissolved, or adsorbed) at any given time from any species across the tree of life.

The analysis of eDNA without considering the existence of these different states has been useful in biodiversity monitoring and conservation applications, but there is a recent shift to consider the states of eDNA to improve knowledge on its persistence and transport in the environment (Mauvisseau et al., 2022; Nagler et al., 2022). Understanding the ecology of eDNA states can also aid in overcoming challenges associated with eDNA analysis such as confirming the current occupancy or relative abundance of surveyed biodiversity (Deiner et al., 2017; Mauvisseau et al., 2022; Nagler et al., 2022). This is because the detection probability of a species’ eDNA in the environment is dependent on its production rate, degradation rate, and transport rate from the source (Barnes & Turner, 2016). For example, a rapid eDNA degradation rate can lead to false negative detection inference for a species’ presence (i.e., the eDNA disappears faster than it can be sampled, but the species is present in the habitat). While a slow eDNA degradation rate can increase persistence and lead to a false positive inference of the species’ presence when in fact it is no longer in the habitat. The degradation rate of eDNA is thus a pivotal parameter to measure and understand its behavior across environmental conditions since the rate change alone can lead to false interpretations of the presence of a species.

The degradation rate of eDNA is hypothesized to be governed by several factors including extracellular nucleases secreted by microorganisms, which are themselves influenced by abiotic conditions like temperature, pH, and light irradiation (Barnes & Turner, 2016; Harrison et al., 2019; Lamb et al., 2022). The rate of degradation has recently been hypothesized to be influenced by the state of eDNA as well (Barnes & Turner, 2016; Harrison et al., 2019; Mauvisseau et al., 2022; Nagler et al., 2022). For instance, membrane-bound DNA may remain protected from extracellular enzymatic degradation, while dissolved DNA may be more susceptible to degradation without the protection of its cellular and organellar membrane (Torti et al., 2015). Similarly, numerous studies demonstrate that adsorbed DNA can remain protected from degradation for hundreds of years (Barrenechea Angeles et al., 2023; Cai et al., 2006b; Capo et al., 2021; Demanèche et al., 2001).

The state of the eDNA has a direct impact also on its transport potential in the environment. For example, eDNA states with different sizes and settling velocities will impact their transport distance (Jo & Yamanaka, 2022; Pont et al., 2018). Pont et al. (2018) found that eDNA in rivers behaves similarly to Fine Particulate Organic Matter (FPOM) and the settling velocity i.e. vertical transfer of eDNA is the primary predictor of eDNA downstream transport distance. But it is also likely that different states of eDNA have different properties affecting their transport (Barnes & Turner, 2016). For example, membrane-bound DNA and adsorbed DNA may exhibit higher settling velocities compared with dissolved DNA, resulting in lower transport potential of these two sources in rivers or higher settling velocities from the surface waters in lentic systems such as lakes (Jo & Yamanaka, 2022). Thus, particle behavior may also be influenced by water body type as well.

Isolation and independent analysis of a chosen eDNA state may be desirable for various applications. For instance, to estimate current occupancy, adsorbed DNA or dissolved DNA may not be fully reliable as these pools may remain protected in their adsorbed state, can resuspend and contribute to the eDNA collected in a water sample (Shogren et al., 2017; Turner et al., 2015). Investigating the current occupancy of species might thus consider a membrane-bound eDNA state as the most appropriate target state. Conversely, an application such as total biodiversity estimation requires high-resolution sampling both temporally and spatially. However, adsorbed DNA pools may represent information on diversity beyond current or seasonal occupancy due to the passive collection and protection of eDNA over time in the adsorbed state (Cai et al., 2006b; Kirtane et al., 2019; Sakata et al., 2020; Turner et al., 2015).

However, the hypotheses that predict the decay and transport behavior of individual eDNA states have not been empirically tested. This is because of the lack of methods to isolate and independently analyze the persistence and transport of individual eDNA states. A small fraction of studies have attempted to separate states of eDNA from a pool of total eDNA for independent analysis of each (Corinaldesi et al., 2005; Lever et al., 2015; Yuan et al., 2019). Corinaldesi et al. (2005), utilized extraction methods to separate microbial extracellular DNA from membrane-bound DNA from the same marine sediment sample. Yuan et al. (2019), separated adsorbed, membrane-bound, and dissolved states of eDNA to investigate the distribution of antimicrobial resistance genes in wastewater. Lever et al. (2015), investigated the performance of various methods and protocols to isolate prokaryotic DNA from different states in the water column, soil, and sediment. All these studies relied on a few key sample processing principles: 1) preventing unintentional cell lysis of membrane-bound DNA; 2) prevention of adsorption of dissolved DNA to particles; 3) causing desorption of adsorbed DNA; 4) size sorting to fractionate dissolved DNA away from adsorbed DNA and membrane-bound DNA, 5) desorption of adsorbed DNA with subsequent separation of this newly desorbed DNA from membrane-bound DNA.

Generally, dissolved DNA is separated from the total eDNA pool via membrane filtration (Figure 1). This involves passing a water sample through a fine pore size filter (usually ~0.2 μm) which should allow dissolved DNA to pass through into the filtrate, while membrane-bound DNA and adsorbed DNA remains on the filter material (Barnes & Turner, 2016; Lever et al., 2015; Sassoubre et al., 2016). Once dissolved DNA is separated using this method into the filtrate, it is concentrated and purified making it suitable for downstream molecular analysis. Ethanol precipitation (Figure 1) is one of the most commonly used methods for extracting dissolved DNA (Lever et al., 2015). The separation of adsorbed DNA from membrane-bound DNA remaining on the filter membrane requires the desorption of adsorbed DNA while minimizing membrane lysis in the process. In the case of adsorbed DNA, the sugar-phosphate backbone is likely covalently bound to hydroxyl groups on particle surfaces such as clay to create chemically adsorbed DNA (Mauvisseau et al., 2022). This can be reversed using phosphate-containing buffers at high pH (Figure 1) (Lever et al., 2015; Mauvisseau et al., 2022; Yuan et al., 2019). Once the formerly adsorbed DNA is desorbed it is expected to go into solution and become dissolved DNA. It can then be separated from the still intact membrane-bound DNA via filtration as before or centrifugation to cause the membrane-bound DNA to form a pellet and the supernatant transferred to remove the newly desorbed DNA. The remaining membrane-bound DNA can then be isolated and purified using membrane lysis and purification steps for downstream molecular analysis (Figure 1) (Lever et al., 2015). Following these methods sequentially suggests that it may be possible to isolate and study the different eDNA states from the same water sample.

**Figure 1:**
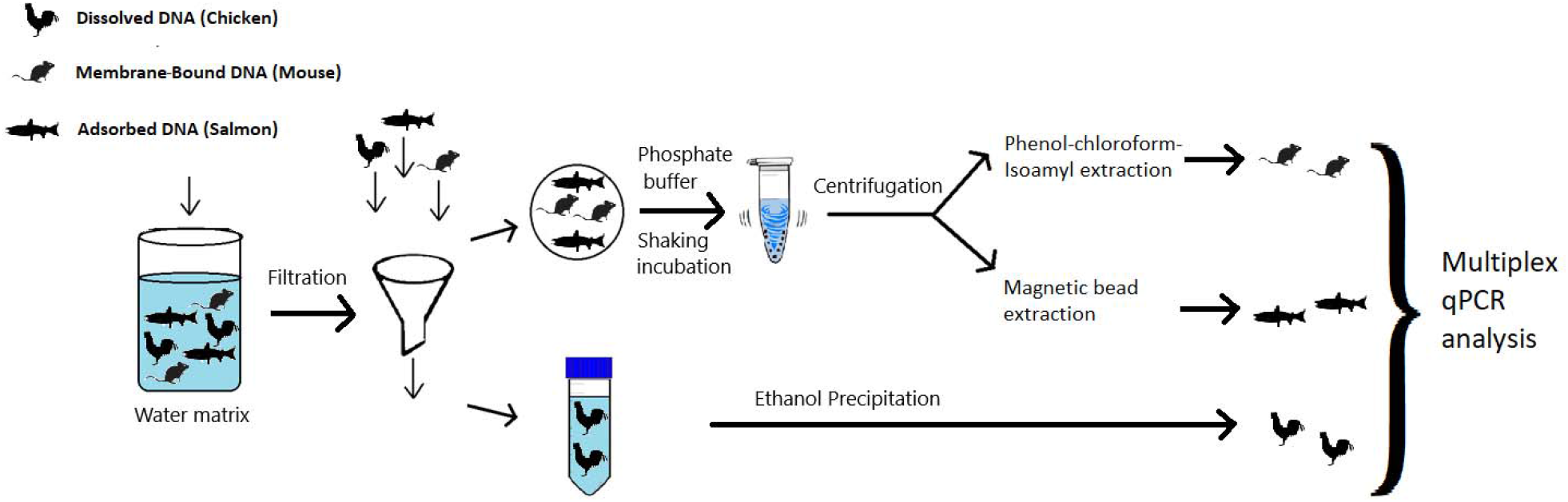
Experimental workflow to isolate eDNA states. Chicken DNA was spiked in the dissolved state, mouse cells were spiked to represent the membrane-bound DNA, and salmon DNA bound to clay was spiked as the adsorbed state.

In this study, we evaluated whether a single protocol can effectively isolate and have a high recovery of different states of eDNA. If successful, this method would result in the ability to separately analyze the community composition measured from each state of eDNA from a single sample. We used species-specific state-controlled spikes where each species represented one state of eDNA (Figure 1). Chicken DNA, mouse cells, and salmon DNA bound to clay particles were used as proxies for dissolved, membrane-bound, and adsorbed DNA states respectively. The separation method consisted of using a 0.22 μm filtration membrane to isolate dissolved DNA from membrane-bound and adsorbed DNA. The filter membrane then was treated with a phosphate buffer to isolate adsorbed DNA from membrane-bound DNA. DNA for each state was recovered by performing ethanol precipitation, phenol-chloroform-isoamyl extraction, and magnetic bead extraction methods to recover dissolved, membrane-bound, and adsorbed states respectively. This experiment permitted the evaluation of two parameters of interest for eDNA state-sorting:1) state-specific isolation and 2) state-specific recovery. State-specific DNA isolation is evaluated based on the presence of a non-target DNA state in a protocol designed to result in a given target state of eDNA. State-specific DNA recovery is used to evaluate the efficiency of a DNA extraction protocol to recover the target eDNA state relative to the spike. The ideal state-specific extraction method should have low contamination from non-target states and high DNA recovery. Furthermore, we investigated the effect of water chemistry on state sorting by replicating the experiment in different water matrices. Lastly, we tested interactions of eDNA states by spiking them independently or all states together.

## 2. Methods

### 2.1 Creation of eDNA states

#### Adsorbed DNA State

Sheared salmon sperm DNA (Invitrogen, Waltham, MA) was diluted to 100 ng/μL in 6 mL nuclease-free molecular grade water (Sigma-Aldrich, St. Louis, MO) in a 15 mL tube with 300 g (50 mg/mL) montmorillonite clay K10 (Fluka, Buchs, CH). One no-adsorbent control tube was created by diluting salmon DNA to 100 ng/μL in 1 mL nuclease-free molecular grade water, but with no clay. The tubes were shaken at 600 rpm for 48 hours. At 48 hours, the tube with the salmon DNA and clay was centrifuged at 4500 xg for five minutes and the supernatant was separated from the pelleted clay with a pipette. The pelleted clay was then washed using 6 mL nuclease-free molecular grade water by vortexing followed by centrifugation at 4500 xg for five minutes and the supernatant was separated to remove any non-adsorbed salmon DNA. This wash process was repeated one more time to remove any non-adsorbed salmon DNA. Finally, 4.5 mL of nuclease-free molecular grade water was added to suspend the clay pellet to create the adsorbed eDNA state spike. The control tube and all the supernatants from each washing step were stored in independent tubes at −20 °C.

#### Dissolved DNA State

DNA from ten (~0.25 g each) pieces of store-bought chicken breast was extracted using the DNeasy Blood and Tissue Kit (Qiagen, Hilden, Germany) according to the manufacturer’s protocol. Each extraction was eluted in 200 μL Buffer AE. The ten extractions were then combined and vortexed to create the dissolved DNA spike.

#### Membrane-bound DNA state

Mouse skin cells from cell line B16-F10 derived from mouse C57BL/6J (Jackson Laboratories, ME, USA) were resuspended in Dulbecco’s Modified Eagle’s Medium (ThermoFisher Scientific, Waltham, MA) containing 10 % Fetal Bovine Serum (ThermoFisher Scientific, MA) and 1 % Penicillin-Streptomycin (10,000 U/mL) (ThermoFisher Scientific, MA) in a 15 mL tube. Cells were spun at 125 x g for 5 minutes and the cell pellet was resuspended in 10 mL growth media and seeded into a tissue culture dish (TPP, Horsforth, UK). Cells were incubated for 10 days at 37 °C, 5 % CO2, and 95 % humidity to let them attach and recover to a concentration of 1×10^6^ cells/m to 20 million cells counted using an automated Cell Counter System (Countess TC20, Biorad, Hercules, CA) which assessed cell viability via trypan blue exclusion. These cells were spun down at 125 x g for 5 minutes in a 50 mL tube and resuspended in 20 mL (1×10^6^ cells/mL) of fresh growth media and used as a spike within 6 hours. The tube was centrifuged at 125 x g for five minutes to pellet the cells. Before spiking the cells, the supernatant was removed using a pipette and discarded. The pellet was then washed to remove any dissolved DNA by resuspending the pellet in 30 mL of Phosphate Buffer Saline (PBS) solution (0.137 M sodium chloride, 0.0027 M potassium chloride, 0.01 M sodium phosphate dibasic, 0.0018 M potassium phosphate monobasic, pH = 7.4). This was followed by centrifuging at 125 x g for five minutes, and the PBS supernatant was discarded as before. The cells were then resuspended in 50 mL of PBS to create the membrane-bound DNA spike.

### 2.2 Experimental procedure for state spiking

A total of three water matrices were used in the experiment: Milli-Q tap water (from Independent Q-POD® ultrapure water dispensing unit (Merck, Darmstadt, Germany), water from Lake Zurich, and water from Sihl River. Ten liters of water near Lake Zurich outlet (47°21’59.2”N 8°32’39.7”E) and Sihl river (47°22’35.8”N 8°32’07.4”E) were collected on October 7, 2021, in the morning of the experiment and transported to the lab within 1 hour. At the lab the pH, turbidity (absorbance), and temperature (°C) of the water matrices were tested using a HI-98194 multiparameter probe (Hanna Instruments, Woonsocket, RI) (Table 1). Three replicates for each water type were created for five treatments (Figure 2). The treatments consisted of spiked DNA from one of each state (i.e., membrane-bound DNA, adsorbed DNA, and dissolved DNA), one where all three states combined were combined, and a control with no spiked DNA (Figure 2). For each treatment, the desired state/s were spiked into 50 mL of water matrix. The volume of spiked states was 500 μL for membrane-bound DNA, 100 μL for adsorbed DNA-bound clay solution, and 50 for of dissolved DNA (Figure 2). The volumes were chosen for ease of spiking and to reduce the chance of accidental double-spiking. The water was then filtered through a 0.22 μm Isopore polycarbonate filter (GTTP02500, Millipore, Burlington, MA) in 25 mm Swinnex filter holders (Millipore, Burlington, MA) using a 50 mL syringe (Figure 1). 15 mL of the filtrate was transferred to a 50 mL falcon tube for dissolved DNA extraction. After filtration, air was passed through to remove any residual water and the filter was immediately removed from the housing and placed in a 1.5 mL tube with 600 μL of phosphate buffer (0.12 M Na_2_HPO_4_, 0.12 M NaH_2_PO_4_, pH = 9) and shaken at 400 rpm for 20 min (Figure 1). The tube was then centrifuged at 13,000 rpm for 2 min. The supernatant was aspirated with a pipette and stored in a separate 1.5 mL tube. The tube with the supernatant was used to extract the adsorbed DNA, while the tube with the filter and pellet was used to extract the membrane-bound DNA. All three fractions (filtrate, supernate, and filter with pellet) were immediately frozen at −20 °C until DNA extraction.

**Figure 2:**
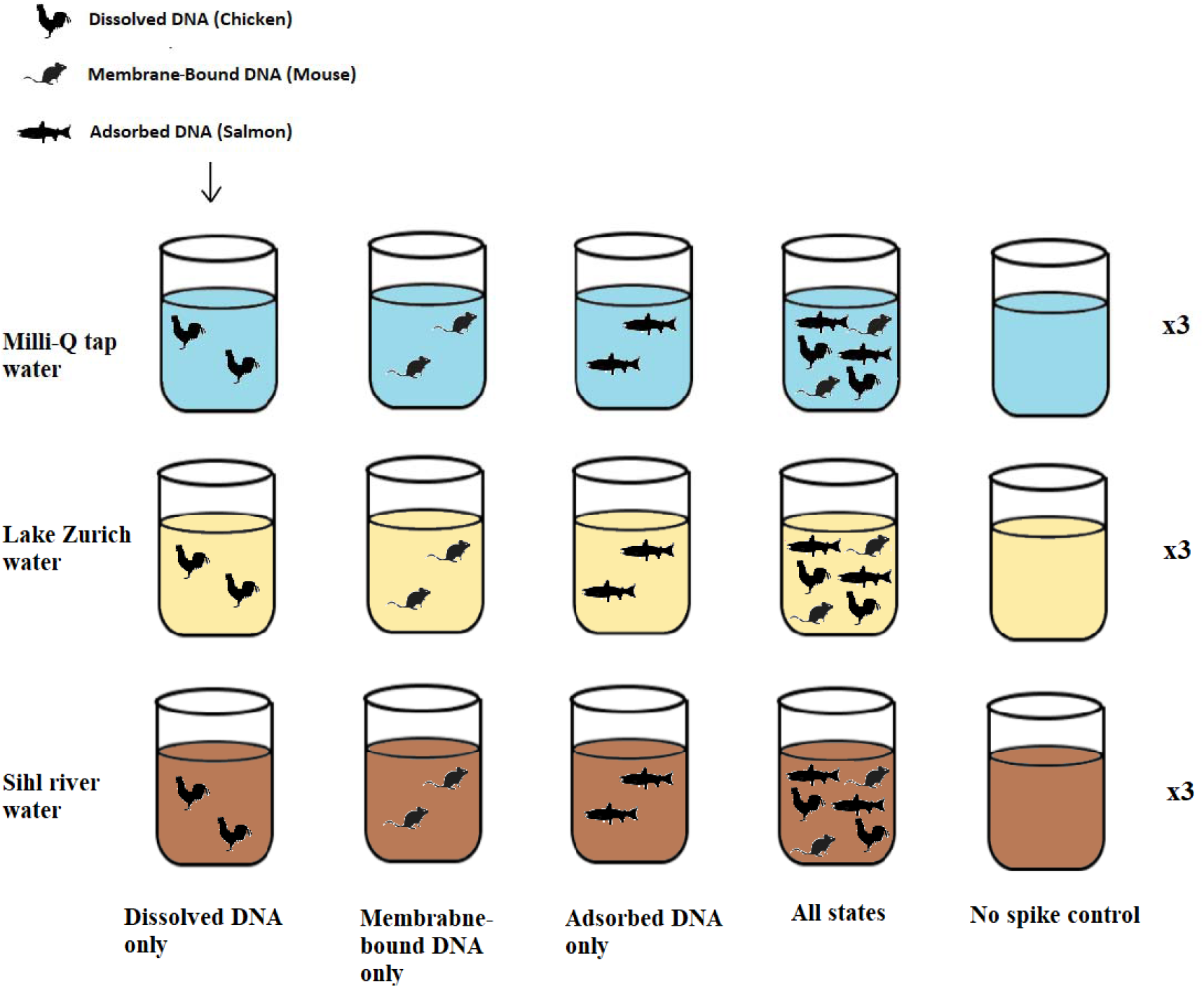
The experimental design used to test the influence of the water matrix and individual vs multiple spiked states on the isolation of eDNA states. All treatments were performed with three replicates each for a total of 45 experimental samples including none-spiked controls.

**Table 1:**
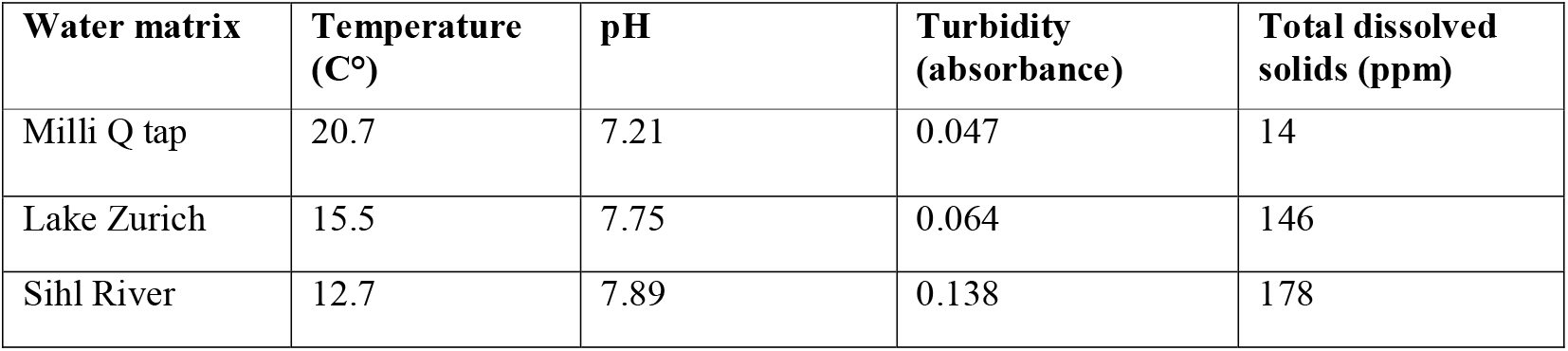
Characteristics of different source water matrices used in this study

### 2.3 DNA extraction methodologies for state separation

The three fractions of the DNA were extracted using three methods chosen specifically to isolate the desired state of eDNA (Figure 1). The dissolved DNA in the filtrate was concentrated using ethanol precipitation, the membrane-bound DNA on the filter and in the pellet was extracted following the lysis step using phenol-chloroform-isoamyl purification and concentrated using ethanol precipitation, and the adsorbed DNA in the phosphate buffer was extracted using a magnetic bead extraction protocol. One negative control was included in every batch of extractions for each method (N = 9).

#### Ethanol precipitation

15 mL of filtrate was used for the isolation of dissolved DNA using ethanol precipitation. Samples were thawed and 1.2 mL of 5M sodium chloride and 33 mL of absolute ethanol (200-proof) were added to the tube. The tube was vortexed and incubated overnight at −20 °C. The tubes were then centrifuged at 10,000 xg at 4 °C for one hour. The supernatant was discarded. 5 mL of 75 % ethanol was added, inverted by hand ten times, and centrifuged at 10,000 xg for 30 mins. The supernatant was discarded, and the pellet was air-dried for 30 minutes. The pellet was then dissolved in 100 μL TE buffer which was then passed through the ZYMO Onestep PCR inhibitor removal kit and stored in 1.5 mL tubes at −20 °C until molecular analysis. This inhibitor removal step was used only for dissolved DNA samples extracted with the ethanol precipitation method.

#### Phenol-chloroform-isoamyl extraction

The membrane-bound DNA from the filters was extracted using a phenol-chloroform-isoamyl (PCI) protocol (Deiner et al., 2015). We added 700 μL of Longmire Lysis Solution (100 mM Tris pH 8.0, 0.5 mM EDTA, 0.2% SDS, 200 mM NaCl) and 12 μL Proteinase K (40 mg/mL) to each of the 2 mL tubes containing the filters. The tubes were gently vortexed prior to overnight incubation at 56 °C to facilitate cell membrane lysis. After the incubation, the lysate was transferred to a new sterile 2 mL tube with a pea-sized volume of grease (high vacuum, Dow Corning®). We then added 550 μL of PCI (25:24:1, Sigma, buffered pH8.0) to all tubes followed by shaking at 20 °C at 1,000 rpm. The tubes were then centrifuged at 10,000 xg for five minutes. The supernatant was transferred to another new sterile 2 mL tube with a pea-sized volume of grease to which we added 550 μL of CI (24:1, Sigma). This tube was also shaken for 5 min at 1000 rpm followed by centrifugation at 10,000x for 5 min. The supernatant was transferred to new 2 mL tubes (without grease) containing 44 μL of 5M NaCl and 1,100 μL of 200-proof ethanol and incubated at −20 °C overnight. The incubated tubes were centrifuged for 30 min at 10,000 x at 4 °C. The supernatant was carefully pipetted out and the pellet was washed twice with 75 % ethanol. The pellet was then allowed to air dry and eluted in 100 μL of TE buffer until molecular analysis.

#### Magnetic bead extraction

A magnetic bead extraction was used to extract and purify formerly adsorbed DNA in phosphate buffer using a version of Powersoil® DNA isolation protocol (Qiagen, Hilden, Germany) using homemade reagents (Sepulveda et al., 2019). The 600 μL of supernatant phosphate buffer containing the desorbed DNA was pipetted into a new 2 mL tube, ensuring the filter or the pellet at the bottom of the tube was not disturbed in the process. We then added 100 μL of protein precipitation solution and inhibitor flocculation solution and vortexed for ten seconds. The tubes were then placed in the freezer at 20 °C for 20 minutes. The tubes were removed and vortexed for ten seconds before centrifugation at 10,000 xg for five minutes. The supernatant was transferred to a new tube with 100 μL of 20% Sera-Mag SpeedBead Carboxylate modified magnetic beads (GE Healthcare Life Sciences, Pittsburgh, PA) in hybridization buffer. The tube was gently mixed by inversion and another 100 μL of hybridization buffer was added. The tube was gently mixed by inversion (10 x) and incubated at room temperature for ten minutes. The tube was then placed on a magnetic rack on a shaker (400 rpm) and shaken until all the beads migrated to the magnet (~ 20 minutes). The supernatant was then pipetted out without disturbing the magnetic beads. Two wash steps were performed where 1 mL of 75 % ethanol was added to the tube. The tube was then removed from the magnetic rack and vortexed for ten seconds, placed back onto the magnetic rack, and shaken until all the beads migrated to the magnet (~ 5 min). The ethanol was then pipetted out without disturbing the magnetic beads. The ethanol wash process was repeated one more time. The tubes were removed from the magnetic rack and air-dried for 20 minutes. The beads were then suspended in 100 μL TE buffer and pipette mixed until in solution and incubated at room temperature for ten minutes. The tube was then placed back onto the magnetic rack for 5 minutes and the TE buffer eluate was pipetted out and passed through a 2 mL EconoSpin® Mini Spin column (Epoch, Fremont, CA) by centrifuging at 10,000 xg for one min to remove any residual magnetic beads in the solution and stored at −20 °C until molecular analysis.

### 2.4 Development of target-specific primers and TaqMan hybridization probes

We designed a multiplex quantitative PCR (qPCR) with four parallel assays to be run on the Roche 480 light cycler (Roche, Basel, Switzerland). Compatible fluorescent dyes (FAM, VIC, TexasRed, and CY5) were selected as recommended by the PrimeTime Multiplex Dye Selection tool (web tool available from IDT DNA). Reference sequences for primer design were obtained from GenBank (Clark, Karsch-Mizrachi, Lipman, Ostell, & Sayers, 2016) (Table S1). To detect and quantify mitochondrial eDNA from mice (*Mus musculus*) and chicken (*Gallus gallus*) we designed TaqMan® qPCR assays targeting the mitochondrial NADH dehydrogenase subunit 2 (ND2) gene, a well-established phylogenetic marker in vertebrates. As a nuclear marker, the single copy gene TGFb1 coding for the transformation growth factor 1 in mice was selected. Previously designed chum salmon (*Oncorhynchus keta*) primers for the cytochrome oxidase I gene (COI) were used with a modified TaqMan® probe that did not have the minor grove binder (Homel et al., 2021) (Table 2) to decrease the cost of the probe. The TaqMan assays for detection and quantification of the nuclear Tgfb1 gene in mice and mitochondrial ND2 genes of chicken and mice were designed using the Primer Express Software version 3.0 (Applied Biosystems, Waltham, MA) using default parameters.

**Table 2:**
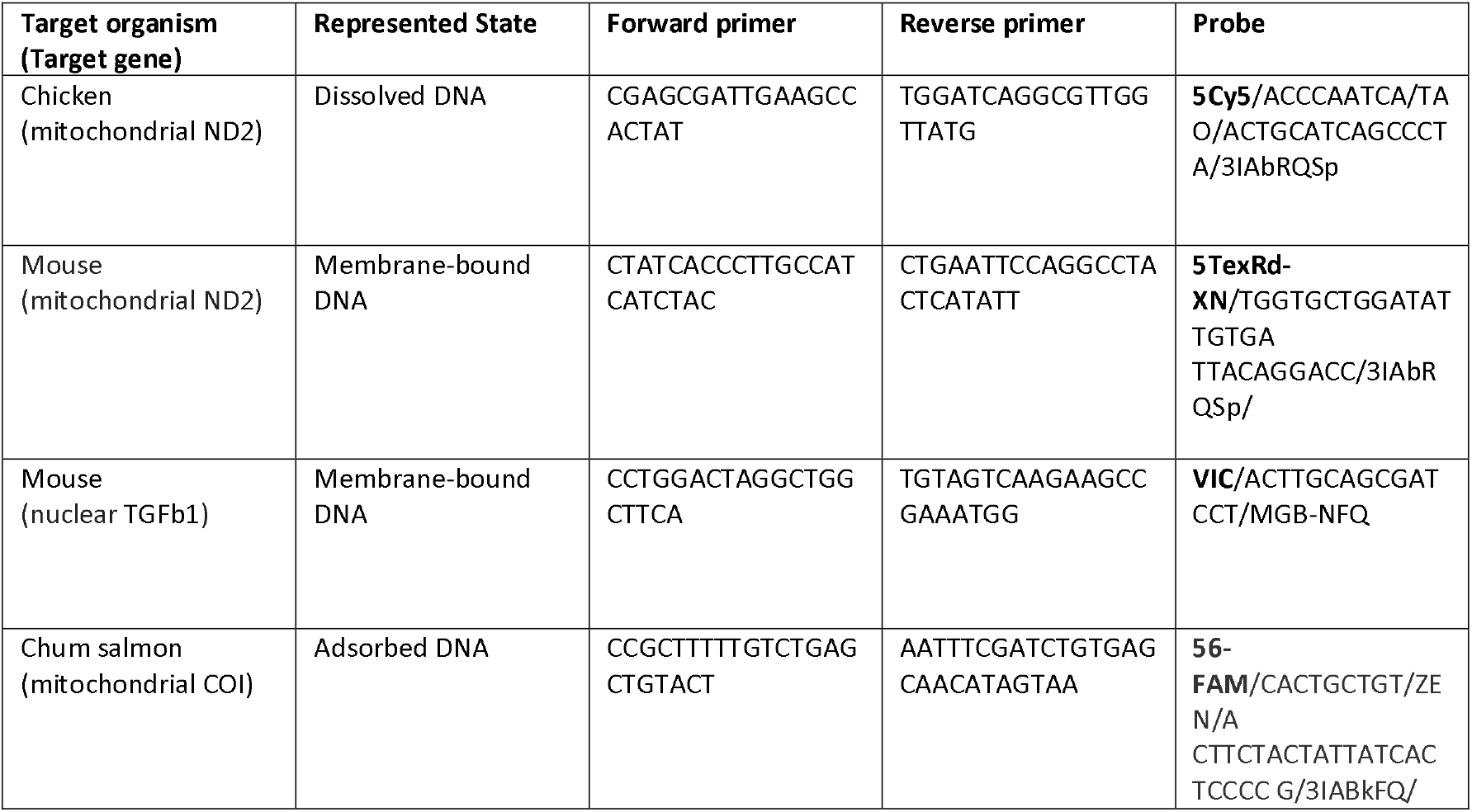
qPCR assays with their corresponding target genes, represented eDNA state and fluorophores (in bold).

#### Specificity testing

All qPCR assays were tested for specificity in-silico and experimentally. The in-silico testing was conducted with NCBI Primer-BLAST tool (Ye et al., 2012) and OligoAnalyzer tool (IDT, Coralville, IA). In Primer-BLAST, the specificity parameters were set to ensure a minimum of three mismatches and at least two mismatches within the last five base pairs of the 3’ end on each primer and probe between the target and non-target organisms used in this study. OligoAnalyzer was used to test the likelihood of dimmer formation between the various primers and probes. Using the default “qPCR” parameters we checked that ΔG > −9 kcal/mole ensured a low likelihood of self- or hetero-dimmers formation in between any primer and probe combinations. To experimentally test the specificity of the multiplex qPCR we amplified standard curve of a single target in the multiplex reaction setup described below. This was repeated by using each of the four target amplicons independently as template in the multiplex qPCR setup. The resulting data was analyzed for cross-amplification or cross-reporting of targets as only one target should be reported from the multiplex qPCR regardless of all assays being available. The efficiencies of the single-species standard curves were compared to the efficiency of the multiplexed standard curves to ensure reliable quantification. The multiplex qPCR negative controls used throughout the experiment ensured no false positives due to dimmer formation.

#### qPCR preparation and cycling conditions

The qPCR reactions were performed in 10 μL reactions in 384 well plates on a Roche Light Cycler 480. Each reaction included 5 μL Taqman™ Multiplex Master Mix (Applied Biosystems™), 0.03 μL of each primer at 100 nM, 0.025 μL of each Taqman probe at 100 nM, 1 μL DNA extract, and 3.92 μL of molecular grade water to bring the volume up to 10 μL. For simplex qPCR, the same reaction mixture was used but only one set of primers and probes were added and the volume of molecular grade water was adjusted to keep the reaction volume to 10 μL. After an initial incubation for ten minutes at 95 °C, we performed 40 cycles with a denaturation step for 15 seconds at 95 °C and an annealing/extension step for 30 seconds at 60 °C. For the preparation of all qPCR plates, we used the mosquito® LV pipetting robot (SPT Labtech Ltd, England) for the efficient and accurate preparation of qPCR plates.

#### qPCR quality control and data interpretation

The Light Cycler was calibrated for multiple emission spectra for the multiplex qPCR using a color compensation protocol utilizing the four fluorophores used in this study. We incorporated six replicates of the six-point standard curve on each qPCR ranging from 10^7 copies/reaction to 100 copies/reaction. These standards were made by combining four individual gBlock gene fragments (Integrated DNA Technologies) that represent the target sequences from the four qPCR assays used in this study (Table S2). The qPCR efficiency was calculated using [E = −1+10^(-1/slope)^] where E is the qPCR efficiency and the slope is calculated with pooled six-point standard curves from all plates for enumerating copy numbers of the target amplicons. This efficiency and intercept were then used in the quantification of our experimental replicates by converting Cp values to copy numbers. We also used this pooled standard curve to determine the Limit of Detection (LOD) and Limit of Quantification (LOQ) using previously described statistical criteria (Klymus et al., 2020). The LOD is described as the lower standard dilution concentration where 95 % of the replicates demonstrate amplification and the LOQ is described as the lowest standard concentration with a coefficient of variation (CV) value below 35 %. Each qPCR plate also included six qPCR negative control wells with molecular grade water instead of template DNA to identify any contamination in the reagents or during qPCR setup.

### 2.5 eDNA state recovery

#### Quantification of total eDNA yield

The total eDNA yield of all treatment samples (N = 45), each spike (N = 3), and extraction controls (N = 9) were measured in 384 well plates using reagents from Qubit dsDNA HS Assay Kit (Qubit Digital, London, UK), and analyzed by Spark® Multimode Microplate Reader (Tecan, Männedorf, CH). We used a seven-point standard curve at concentrations of 0, 0.1, 0.2 0.5, 1, 5, and 10 ng/μL by diluting the 10 ng/μL standard provided by the manufacturer. Each reaction well consisted of 48 μL of Qubit HS 1x reaction mixture and 2 μL of DNA standard, sample, spike, or control. Accurate pipetting was facilitated by a mosquito® LV pipetting robot (SPT Labtech Ltd, England). All standards and samples were analyzed in triplicate.

#### State-specific isolation efficiency and percent recovery

To calculate the extraction recovery of the DNA added to each replicate of the experiment, first, the spiked DNA for each of the three states was quantified. The concentration of target DNA recovered from the experimental samples was then quantified and compared with the spike to calculate the percent recovery of each DNA state.

For quantification of the dissolved state spike, 1 μL of the dissolved DNA spike was directly analyzed in triplicate using simplex qPCR to enumerate the dissolved DNA target copies per uL. This value was multiplied by the spike volume (50 μL) to calculate the total copy number of the spiked DNA. Using Equation 1, percent recovery was calculated where C_s_ is the number of DNA copies spiked in the dissolved state (per 15 mL), C_i_ is the initial spike concentration, Vtot is the total volume of the filtrate (50 mL), and Vs is the volume of the filtrate analyzed (15 mL). This volume correction is necessary as only 15 mL of filtrate could be used in the isolation id dissolved DNA using the ethanol precipitation method.

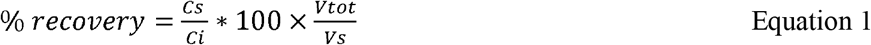

For quantification of the membrane-bound DNA spike, 2 mL of mouse cell spike solution was frozen at −20 °C. Three replicates of the spike were created by extracting 50 μL of the saved spike using the same phenol-chloroform-isoamyl protocol described above and analyzed with qPCR to enumerate the membrane-bound DNA target copies spiked into the experiments.

To calculate the percent recovery of the membrane-bound DNA from experimental samples we used equation 2, where C_s_ is the concentration of mouse DNA (mitochondrial or nuclear) detected in experimental samples (per 50 mL) and C_i_ is the initial spike concentration of mouse DNA.

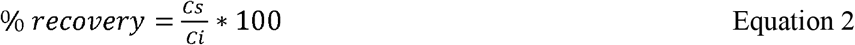

The concentration of adsorbed DNA spike was calculated using equation 3, where C_sp_ is the concentration of spiked adsorbed DNA (copies DNA/mg clay), C_i_ (copies/μL) is the initial concentration of salmon DNA solution, V_i_ (μL) is the initial volume of salmon DNA solution, C_sup_ (copies/μL) is the concentration of salmon DNA in the supernatant after 48 h of adsorption, C_w1_ (copies/μL) and C_w2_ (copies/μL) are concentrations of salmon DNA in the supernatant of the first and second wash step, V_w_ (μL) is the volume of water used in the wash steps and M_c_ (mg) is the mass of montmorillonite clay added to the adsorption reaction. Finally, the adsorbed DNA was suspended in 3 mL of nuclease-free molecular grade water to achieve a final adsorbed DNA spike concentration of 100 mg/mL. Concentrations of target DNA copies in C_i_, C_sup_, C_w1_, and C_w2_ were quantified in triplicate using simplex qPCR.

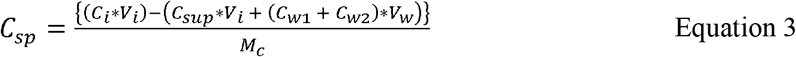

To calculate the extraction recovery of the adsorbed DNA in each sample, equation 4 was used where C_s_ is the concentration of salmon DNA detected from experimental samples (per 50 mL) and C_sp_ is the theoretical adsorbed DNA spike concentration calculated in equation 3.

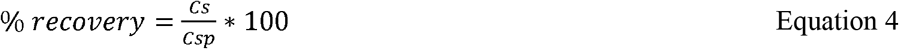

### 2.6 Statistical analyses

We conducted an analysis of variance (One-way ANOVA) using the extraction methods, water matrix type, and the state of spiked DNA as dependent variables, and percent recovery as the independent variable. We used a one-way ANOVA to test whether a state isolation protocol was able to enrich the target state. This test was repeated for each of the three state isolation protocols used. A one-way ANOVA test was also used to test if any state isolation protocol was able to outperform others with respect to percent recovery of the target state. Finally, another one-way ANOVA was also used to determine whether a given water matrix had a significant impact in determining the success of eDNA state isolation based on the increased recovery of a target state. We used a student’s t-test to evaluate the effect of spiking a given state individually in a sample or multiple states spiked together. The t-test was also used for testing the variation in the recovery of mitochondrial and nuclear DNA recovery from spiked mouse cells. We conducted the Shapiro-Wilk test of normality, and Levene’s test to check the homogeneity of variances to ensure our data met the assumptions of parametric t-tests and ANOVA. All ANOVA tests that rejected the null (α = 0.01) were followed up with Tukey’s post hoc test to identify what dependent variables caused a significant difference. All data analysis was conducted in R version 4.1.3 using the package tidyverse package (Wickham et al., 2019).

## 3. Results

### 3.1 Performance of multiplex qPCR assays

None of the qPCR assays used in this study cross-amplified other targets in multiplex reactions. This was confirmed by the lack of non-specific amplification or fluorescence when single target standard curves were added to the multiplex reaction mix. All four multiplex qPCR assays used in the study had a Limit of Detection (LOD) at 10 copies/reaction and the Limit of Quantification (LOQ) was 100 copies/reaction for all targets. The efficiencies of pooled multiplex standard curves from all plates used in the experiment were 0.85, 0.89, 0.85, and 0.93 for salmon (adsorbed DNA), chicken (dissolved DNA), and mouse mitochondrial target (membrane-bound DNA), and mouse nuclear (membrane-bound DNA) assays respectively (Figure S1). This efficiency was comparable with simplex standard curves allowing accurate quantification of the target DNA (Figure S2).

None of the negative controls, including no spike controls, extractions negatives, and qPCR negative controls showed amplification over the LOD in all three qPCR replicates for any of the four targets. However, below LOD concentrations (i.e. < 10 copies/reaction) of target DNA were detected in some no-spike controls. This was observed in one, nine, four and thirteen qPCR replicates for mouse nuclear, mouse mitochondrial, salmon, and chicken targets respectively of a total of 81 no-spike controls qPCR replicates. This was only observed in the no-spike controls processed using methods targeting membrane-bound and adsorbed eDNA. Since the qPCR and extraction negative controls showed no amplification, the cause of this contamination can be incomplete sterilization of beakers between experiments, and/or the natural presence of target DNA from environmental waters from Lake Zurich and the Sihl river. The low concentrations and the nature of this contamination are unlikely to have affected the results of this experiment.

### 3.2 State-specific DNA isolation

Species-specific isolation of DNA was evaluated based on the presence of non-target states of eDNA in a given extraction protocol. Thus, this analysis can be conducted only on treatments where all eDNA states were spiked altogether. None of the protocols tested were able to completely isolate the target eDNA state. Specifically, all states were detected in replicates where they were not expected (Table 3, Figure 3). However, some protocols resulted in limiting non-target extraction and enriching the target eDNA state. For instance, the protocol designed to isolate membrane-bound eDNA (filtration and desorption followed by PCI extraction on the pellet) resulted in increased enrichment of membrane-bound DNA state (One-way ANOVA, F (3, 102) = [152.8], p = 2e-16) as the percent recovery of membrane-bound DNA was significantly higher than that of dissolved (Tukey HSD, p = 0.00, 95% C.I. = [31.46, 43.84]) and adsorbed DNA (Tukey HSD, p = 0.00, 95% C.I. = [31.19, 43.58]) (Figure 3C, D [PCI extraction]). This result was not significantly different between mitochondrial and nuclear targets of membrane-bound DNA (Tukey HSD, p = 0.65, 95% C.I. = [−9.14, 3.45]) (Figure 3C, D, Table 3 [PCI extraction]). In the protocol designed to isolate dissolved DNA, the DNA from both membrane-bound and adsorbed states were detected with similar percent recoveries as dissolved DNA (Table 3, Figure 3 B,C [Ethanol precipitation]). Filtration followed by ethanol precipitation on the filtrate, therefore, did not lead to effective isolation of dissolved DNA from the other two as the percent recovery of dissolved DNA was not significantly higher than that of the adsorbed (One-way ANOVA, F(2, 69) = [5.71], p = 0.005; Tukey HSD, p = 0.12, 95% C.I. = [−0.02, 0.28]) or membrane-bound DNA Tukey HSD, p = 0.29, 95% C.I. = [−0.27, 0.60]). Similarly, the protocol designed for adsorbed state isolation (filtration followed by desorption and magnetic bead extraction on the supernatant) did not isolate adsorbed DNA as the percent recovery of adsorbed DNA was significantly lower than that of membrane-bound DNA (One-way ANOVA, F(3, 97) = [40.82], p = 2.0^-16^; Tukey HSD, p = 0.0, 95% C.I. = (4.20, 9.53). Additionally, the resulting percent recovery of adsorbed DNA was not significantly higher than that of dissolved DNA when processed using the protocol for adsorbed DNA isolation (Tukey HSD, p = 0.56, 95% C.I. = [−1.30, 3.98]) (Table 4, Figure 3A, B [magnetic bead extraction]).

**Table 3:**
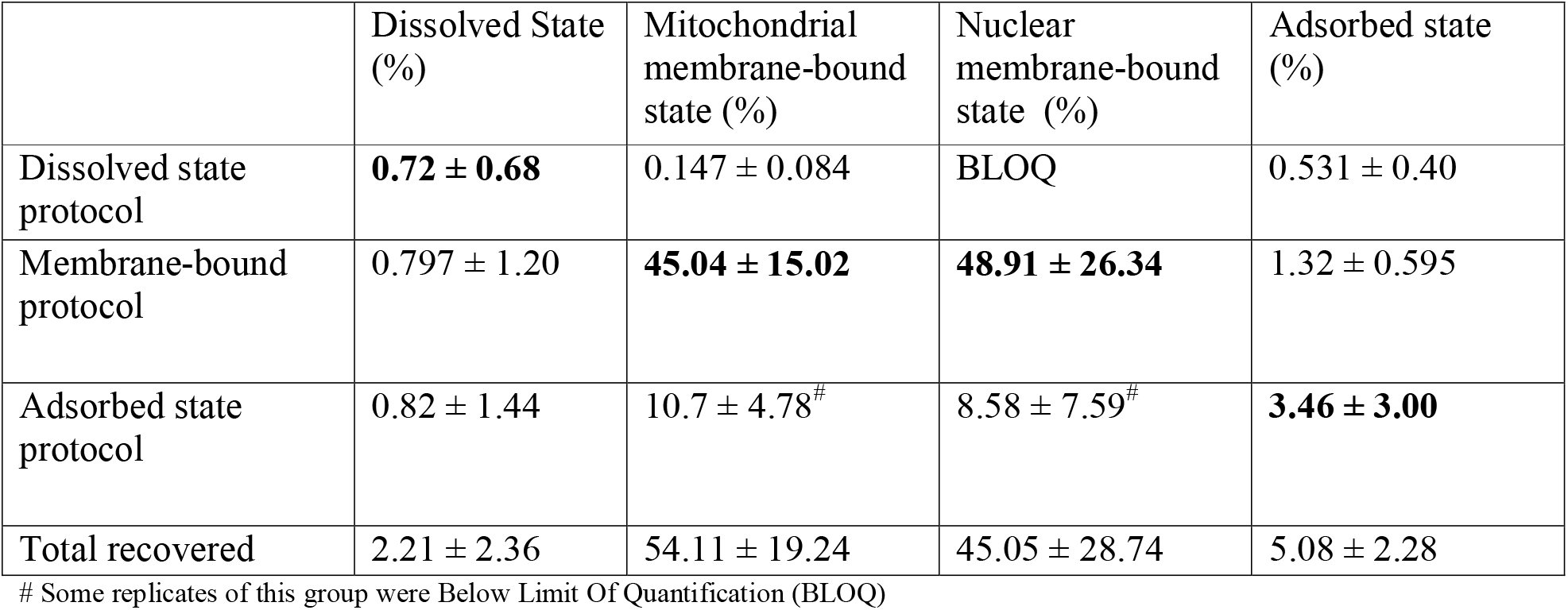
Percent recovery of target and non-target eDNA expressed based on DNA state (columns) and extraction protocol (rows) used to isolate that expected state. Cells in BOLD indicate replicates with high expected recovery of the target eDNA state.

**Figure 3:**
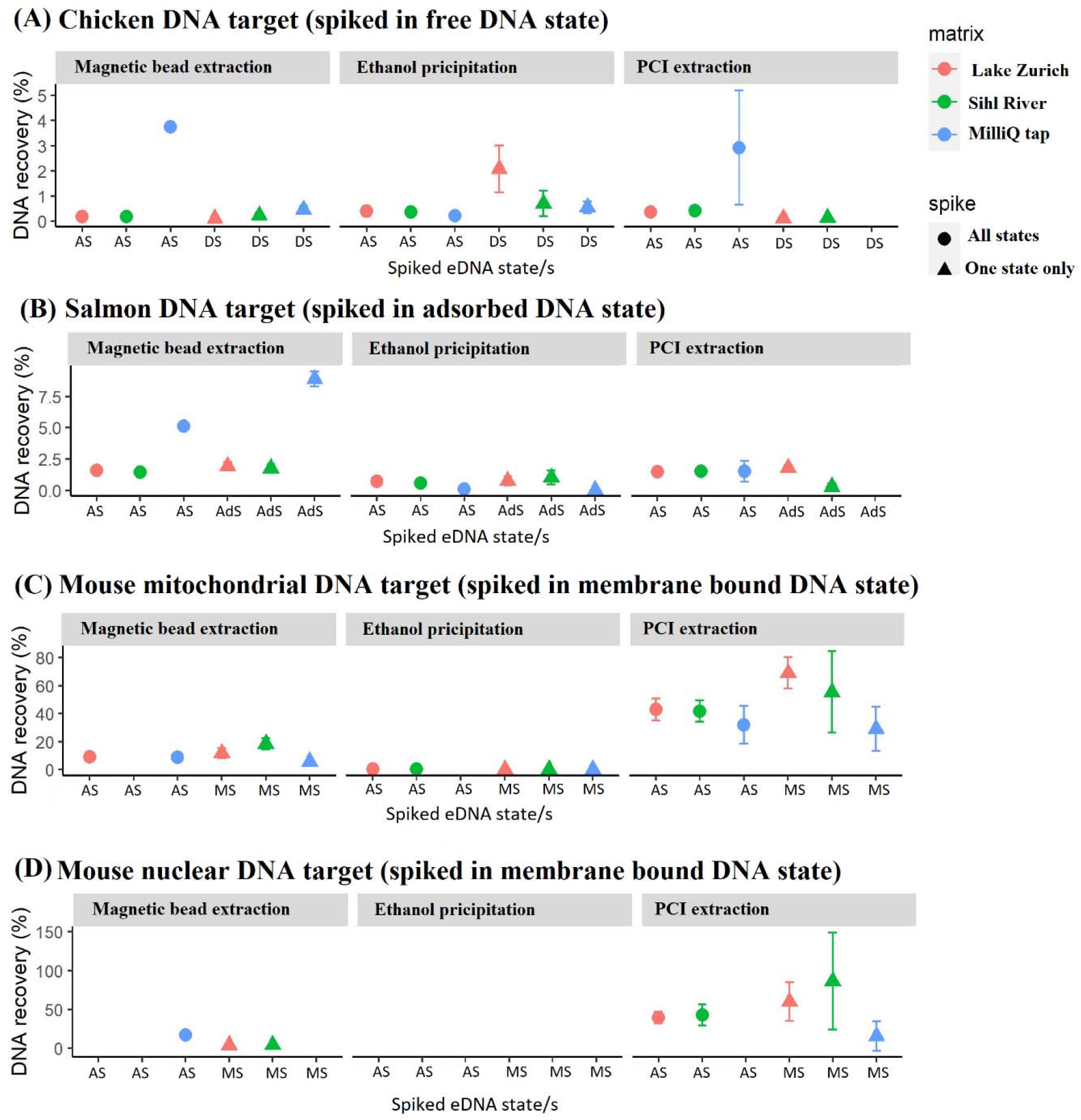
Percent recovery of spiked eDNA states observed in three different extraction methods. Colors represent the water matrix type while the shape of the data points indicates whether all states were spiked together, or the target state spiked by itself. The x-axis shows the spiked eDNA state (AS = All States, DS = Dissolved State, AdS = Adsorbed State, and MS = Membrane-bound State). Error bars indicate the standard deviation between three biological replicates.

### 3.3 State-specific DNA recovery

The state-specific DNA recovery is used to evaluate how much of the target eDNA state was recovered from an experimental unit (i.e. eDNA extraction protocols) irrespective of the presence of non-target eDNA states. The state-specific spike concentrations, in theory, reflect 100 % recovery of a given state (Table 4). The recovery of membrane-bound DNA using the filtration followed by desorption and performing PCI extraction on the pellet was significantly greater than the recovery of membrane-bound state in other methods. This was the case for both mitochondrial (One-way ANOVA, F(2, 149) = [178.4] p = 2e-16) and the nuclear targets of membrane-bound DNA (One-way ANOVA, F(2, 96) = [46.7] p = 7.7e-10) when compared to other isolation protocols (Table 4). There was no significant difference in the percent recovery of mitochondrial and nuclear targets using the membrane-bound isolation protocol (t-test, df = 78.70, t = −0.06, p = 0.94), however, the concentration of mitochondrial marker was more than two orders of magnitude higher in both, the spike and the recovered DNA (Table 4, S3). The recovery of adsorbed DNA was significantly higher using the protocol specially designed for it (i.e. filtration, followed by desorption and magnetic bead extraction on the supernatant) when compared to the other two methods (One-way ANOVA, F(2, 156) = [43.8] p = 8.2e-16). Tukey’s post hoc test revealed significant increases in adsorbed DNA recovery between the magnetic bead method, and both PCI (Tukey HSD, p < 0.01, 95% C.I. = [−3.69, −2.15]) and EtOH precipitation methods (Tukey HSD, p < 0.01, 95% C.I. = [−3.03, −1.46]). The DNA recovery of dissolved DNA was not significantly greater using the protocol designed for isolating dissolved DNA as compared to other protocols (One-way ANOVA, F(2, 150) = [0.12] p = 0.90).

The eDNA yield of the full-process negative controls from the lake and river sample i.e., without any spikes can indicate the magnitude of genetic information available in a given state and further elucidate on total eDNA recovery from each method. The eDNA yields of no-spike controls using the adsorbed state and membrane-bound state protocols were 0.34 ± 0.22 ng/μL and 0.74 ± 0.06 ng/μL respectively for Lake Zurich water, and 0.24 ± 0.25 ng/μL and 1.07 ± 0.36 ng/μL respectively for Sihl river water. The yield of the dissolved eDNA fraction represented by the ethanol precipitation method was below the limit of detection for all water matrices.

**Table 4:** Spike, recovery, and loss of DNA based on the state the DNA was spiked in.

|  | Dissolved State | Membrane-bound state (mitochondrial) | Membrane-bound state (nuclear) | Adsorbed state |
| --- | --- | --- | --- | --- |
| DNA spike concentration (copies/50 ml) | $4.0 \times 10^8$ | $8.5 \times 10^6$ | $3.0 \times 10^4$ | $2.5 \times 10^7$ |
| Total recovery (copies/50 ml) | $8.7 \times 10^6 \pm 9.3 \times 10^6$ | $4.6 \times 10^6 \pm 1.6 \times 10^6$ | $1.4 \times 10^4 \pm 8.6 \times 10^3$ | $1.3 \times 10^6 \pm 5.7 \times 10^5$ |
| Percent recovery (%) | $2.21 \pm 2.36$ | $54.11 \pm 19.24$ | $45.05 \pm 28.74$ | $5.08 \pm 2.28$ |
| Percent DNA lost (%) | $97.78 \pm 2.35$ | $45.88 \pm 19.04$ | $54.95 \pm 28.74$ | $94.92 \pm 2.32$ |

### 3.4 Effect of water matrix and spiking multiple states altogether

Overall, the percent recovery of DNA was not significantly influenced by whether a given state-controlled spike was spiked individually with other eDNA states (t-test, df = 591.82, t = 0.32, p = 0.75). Similarly, the overall percent recovery was not significantly affected by the water matrix (MilliQ tap, Lake Zurich, or Sihl river) they were spiked in (One-way ANOVA, F(2, 276) = [0.05,] p = 0.94). However, in select scenarios, water matrix type did have a significant impact on the recovery of adsorbed and dissolved states of DNA. The recovery of adsorbed DNA, using the magnetic bead extraction protocol, was significantly higher in the Milli-Q water matrix when compared to the recovery in Lake Zurich (One-way ANOVA, F(2, 51) = [112.2], p < 2e-16; Tukey HSD, p < 0.01, 95% C.I. = [4.30, 6.20]) and Sihl river water (Tukey HSD, p < 0.01, 95% C.I. = [4.45, 6.36]), while the recovery of adsorbed DNA was not significantly different in the two environmental waters (Tukey HSD, p < 0.01, 95% C.I. = [−1.02, 0.79]) (Figure 3B [magnetic bead extraction]). Similarly, the recovery of dissolved DNA, using the ethanol precipitation protocol, was significantly higher in the Milli-Q water matrix when compared to the recovery in Lake Zurich (One-way ANOVA, F(2, 51) = [8.06], p = 9.1e-4; Tukey HSD, p < 0.01, 95% C.I. = [4.30, 6.20]) and Sihl river water (Tukey HSD, p < 0.01, 95% C.I. = [4.45, 6.36]), while the recovery of dissolved DNA in the two environmental waters was not significantly different Tukey HSD, p < 92, 95% C.I. = [−1.12, 0.80]) (Figure 3A [magnetic bead extraction]) . Contrary to this pattern, the recovery of membrane-bound DNA, using the PCI extraction protocol, was significantly higher in Lake One-way ANOVA, F(2, 51) = [9.75], p = 2.6e-6; Tukey HSD, p < 0.01, 95% C.I. = [−39.84, −11.17]) Zurich and Sihl river (Tukey HSD, p < 0.01, 95% C.I. = [−32.43, −3.76]) waters compared to Milli-Q water, although the membrane-bound DNA recovery was not significantly different between the two environmental water matrices (Tukey HSD, p = 0.43, 95% C.I. = [−21.74, 6.93]) (Figure 3C, D [PCI extraction]).

In select scenarios, we also observed an increase in the percent recovery of dissolved DNA in non-target protocols, i.e. protocols designed for adsorbed and membrane-bound DNA isolation. The percent recovery of dissolved DNA using the ethanol precipitation protocol (0.216 ± 0.108 %) increased dramatically using the adsorbed DNA protocol (3.76 ± 0.179 %) and membrane-bound DNA protocol (3.93 ± 2.27 %) in Milli-Q tap water when all three states were spiked together (Figure 3A, S3A). This represents an increase in recovery of ~1640% and ~1256 % respectively of dissolved DNA in using protocols targeting the adsorbed and membrane-bound DNA respectively (Figure 3A, S3A). Interestingly, this was the case only when other states, i.e., cells and clay, were present and only in the Milli-Q tap water matrix (Figure 3A, S3A). The treatment with only dissolved DNA spiked in Milli-Q tap water did not show this dramatic increase (Figure 3A, S3A).

## 4. Discussion

Answering questions regarding the ecology of eDNA such as persistence and transport require effective and consistently replicable methods to capture and isolate particular states of eDNA. Here we demonstrate the utility of controlled experiments using state-specific spikes from different species to evaluate DNA extraction protocols, their effect on the isolation and recovery of each of the DNA states to aid in the understanding of the ecology of eDNA in different water chemistries. We show that while no methods were able to isolate a state in its entirety, we could enrich the recovery of each state to some extent. Because of the novel experimental design we also measured and identified the fate of each DNA state when it was not captured using the targeted protocol. The protocols for state sorting were able to enrich the target state, especially in the case of membrane-bound and adsorbed DNA states.

However, there was a significant cross-over between states indicating inefficiencies in sample processing methods and potential dynamics of eDNA states after the collection of the water sample. Additionally, a large proportion of DNA in all states was lost and not recovered by any of the treatments. Further, experimentation using a multiple-state-specific spike model designed here can shed light on the state dynamics of eDNA during the isolation process and help to optimize state isolation protocols.

### Isolation of target states

The results of this study highlight current limitations with state-specific DNA isolation protocols and help identify steps in their optimization. Previous studies have utilized methods for sorting and isolating eDNA states but have not been able to verify their success in doing so (Corinaldesi et al., 2005; Lever et al., 2015; Torti et al., 2015; Yuan et al., 2019). Unverified methods with unknown levels of inefficiencies can lead to misinterpretation of results. Here we show that none of the isolation methods used in these previous studies were able to completely isolate the target state of eDNA, but some were able to enrich a target state. For example, when all three states were spiked together, membrane-bound DNA accounted for a majority (~ 85.6%) of the DNA recovered using a protocol designed for membrane-bound DNA isolation, the adsorbed DNA accounted for two-thirds (~ 68.1 %) of total recovered DNA using the protocol designed for the isolation of adsorbed DNA, however, the dissolved DNA state only accounted for a third (~ 32.5%) of the total recovered DNA from the protocol designed to isolate the dissolved state of eDNA.

### Effect of water matrix characteristics on eDNA state isolation

This study performed all experiments in three water matrices, two environmental (Lake Zurich and Sihl river) and one artificial (Milli-Q tap). Overall, the change in environmental waters did not significantly impact the results of state isolation even though they had different abiotic conditions (Table 1). The least influence was noted in the case of isolation of membrane-bound DNA, probably because the isolation of membrane-bound DNA from other states is primarily a physical separation while other states of eDNA might experience more chemical interactions influenced by water matrix during their separation. For instance, experiments in Milli-Q water matrix led to increased recovery of adsorbed and dissolved DNA in select scenarios compared to the two environmental water matrices.

The recovery of adsorbed DNA was significantly higher in Milli-Q tap water than in the two environmental waters using the adsorbed state protocol (Figure 3). The difference between the Milli-Q tap and environmental waters was the more circumneutral pH, and absence of other particles and organics in the Milli-Q water (Table 1). The phosphates in the desorption buffer may have competitively interacted with these other particles reducing the desorption efficiency and thus the percent recovery. Improving the recovery of the adsorbed state of eDNA requires an improved understanding of the mechanisms that create the adsorbed state of eDNA in the first place. Adsorption of DNA onto mineral surfaces is governed by interactions of multiple mechanisms including electrostatic forces, hydrogen bonding, ligand exchange, and cation bridging (Franchi et al., 1999; Pietramellara et al., 2001; Saeki et al., 2010; Yu et al., 2013). Furthermore, the water chemistry can impact the adsorption mechanisms even when the adsorbent and adsorbate are consistent (Kirtane et al., 2020). pH and ionic strength have been categorized as the driving characteristics of a solution to influence the adsorption mechanisms (Yu et al., 2013). Increased pH (>5) reduces the protonation of DNA bases giving it a net negative charge, thus reducing adsorption via electrostatic forces with predominantly negatively changed mineral clay surfaces, and increasing the effect of cation bridging (Xu et al., 2003; Yu et al., 2013). Increased concentrations of cations in the water matrix increase the adsorption of DNA onto mineral surfaces via cation bridging (Cai et al., 2006a; Levy-Booth et al., 2007).

Dissolved DNA recovery increased by over an order of magnitude when extracted using protocols for membrane-bound and adsorbed DNA extraction but only in one scenario containing the Milli-Q water matrix with all states spiked together (Figure 3). The circumneutral pH of the Milli-Q water likely caused the spiked dissolved DNA to rapidly adsorb to the clay particles and cells in the water leading to an increased recovery of dissolved DNA in method treatments targeted toward the extraction of membrane-bound and adsorbed states (Mauvisseau et al., 2022). This effect required both Milli-Q water and the presence of adsorbents in the water as this increase was not observed in the treatments with environmental water matrices or in Milli-Q water with only dissolved DNA spiked into it. Thus, we recommend the use of synthetic water matrices instead of Milli-Q water in future studies for reproducible controlled experiments that better reflect the water chemistries in the environment.

### Strategies for improving eDNA state isolation and recovery

Due to the novel ability to quantify state-specific extraction efficiencies and the level of isolation of a target state, this experiment also aided to identify various opportunities to improve state sorting and extraction methods. The biggest room for improvement was in the case of extraction of dissolved and adsorbed DNA. This is also intuitive as most development of methods has been inadvertently targeted toward membrane-bound DNA (Pawlowski et al., 2021; Tsuji et al., 2019). The challenge with the extraction of dissolved DNA is that of concentration or aggregation. Unlike the other states, dissolved DNA cannot be easily concentrated via filtration. We utilized ethanol precipitation, the most popular method for dissolved DNA concentration, but alternative methods for aggregation such as column chromatography, magnetic bead extraction, and lyophilization can be tested to improve the recovery of dissolved DNA (Calderón-Franco et al., 2021; Rees et al., 2014; Yuan et al., 2019).

The recovery of adsorbed DNA will be improved by understanding the mechanistic interactions between the particles and DNA. In this case, we used a model mineral clay, montmorillonite, which binds to the sugar-phosphate backbone of DNA via electrostatic attraction or cation-bridging (Saeki et al., 2010; Sheng et al., 2019). Thus, the addition of phosphates to the mix is likely to weaken and replace those bonds, thus desorbing the DNA into solution. In this experiment, we used two phosphates (Na2HPO4 and NaH2PO4) at 0.12M each (Yuan et al., 2019). Other studies have hypothesized hexaphosphates or deoxyribose triphosphates to improve the desorption ability (Direito et al., 2012; Lever et al., 2015). Since the recovery of adsorbed DNA was reduced in environmental matrices as compared with Milli-Q water when treated with the adsorbed state protocol, we hypothesize that the reduction of recovery is attributed to incomplete desorption of adsorbed DNA due to competitive interactions with other particles and organics in the environmental water matrices. Future studies should evaluate the effect of increasing phosphate concentrations and using varied forms of phosphates discussed on the desorption of DNA from complex environmental matrices. In natural waters, DNA is likely to be adsorbed to or otherwise interacting with numerous types of particles such as clays, porous carbons, organic molecules, metal oxides, and even biofilms, etc. (Kirtane et al., 2020; Saeki et al., 2010; Sheng et al., 2019; Sodnikar et al., 2021). As discussed above, the water chemistry impacts the adsorption mechanism and thus the success of desorption strategies. In this experiment, the adsorbed state spike was created by adding montmorillonite clay and salmon DNA in molecular grade water. Hence, future studies should consider using a mixture of complex adsorbents instead of a single model mineral clay and create the adsorbed DNA state spike in relevant environmental or synthetic waters to better replicate the “real-world” behavior of adsorbed DNA state and optimize the methods to isolate it.

## Acknowledgments

We would like to thank Myriam Gwerder, Lukas Sommer, and Benjamin Loos for culturing mouse cells for the experiments. We thank Pascal Opiasa and Killian Zurita de Higes for collecting water matrices and assisting with the experiment. We thank Silvia Kobel for assistance with the Mosquito liquid handler. Data produced and analyzed in this paper were generated in collaboration with the Genetic Diversity Centre (GDC), ETH Zurich. This work and all co-authors have been supported by the European Research Council (ERC) under the European Union’s Horizon 2020 research and innovation program (Grant Agreement No. 852621).

## References

Barnes, M. A., & Turner, C. R. (2016). The ecology of environmental DNA and implications for conservation genetics. Conservation Genetics, 17(1), 1–17. https://doi.org/10.1007/s10592-015-0775-4

Barrenechea Angeles, I., Romero-Martínez, M. L., Cavaliere, M., Varrella, S., Francescangeli, F., Piredda, R., Mazzocchi, M. G., Montresor, M., Schirone, A., Delbono, I., Margiotta, F., Corinaldesi, C., Chiavarini, S., Montereali, M. R., Rimauro, J., Parrella, L., Musco, L., Dell’Anno, A., Tangherlini, M., … Frontalini, F. (2023). Encapsulated in sediments: EDNA deciphers the ecosystem history of one of the most polluted European marine sites. Environment International, 107738. https://doi.org/10.1016/j.envint.2023.107738

Cai, P., Huang, Q., & Zhang, X. (2006a). Microcalorimetric studies of the effects of MgCl2 concentrations and pH on the adsorption of DNA on montmorillonite, kaolinite and goethite. Applied Clay Science, 32(1–2), 147–152.

Cai, P., Huang, Q.-Y., & Zhang, X.-W. (2006b). Interactions of DNA with clay minerals and soil colloidal particles and protection against degradation by DNase. Environmental Science & Technology, 40(9), 2971–2976.

Calderón-Franco, D., van Loosdrecht, M. C., Abeel, T., & Weissbrodt, D. G. (2021). Free-floating extracellular DNA: Systematic profiling of mobile genetic elements and antibiotic resistance from wastewater. Water Research, 189, 116592.

Capo, E., Giguet-Covex, C., Rouillard, A., Nota, K., Heintzman, P. D., Vuillemin, A., Ariztegui, D., Arnaud, F., Belle, S., & Bertilsson, S. (2021). Lake sedimentary DNA research on past terrestrial and aquatic biodiversity: Overview and recommendations. Quaternary, 4(1), 6.

Corinaldesi, C., Danovaro, R., & Dell’Anno, A. (2005). Simultaneous recovery of extracellular and intracellular DNA suitable for molecular studies from marine sediments. Applied and Environmental Microbiology, 71(1), 46–50.

Deiner, K., Bik, H. M., Mächler, E., Seymour, M., Lacoursière-Roussel, A., Altermatt, F., Creer, S., Bista, I., Lodge, D. M., de Vere, N., Pfrender, M. E., & Bernatchez, L. (2017). Environmental DNA metabarcoding: Transforming how we survey animal and plant communities. Molecular Ecology, 26(21), 5872–5895. https://doi.org/10.1111/mec.14350

Deiner, K., Walser, J.-C., Mächler, E., & Altermatt, F. (2015). Choice of capture and extraction methods affect detection of freshwater biodiversity from environmental DNA. Biological Conservation, 183, 53–63. https://doi.org/10.1016/j.biocon.2014.11.018

Demanèche, S., Jocteur-Monrozier, L., Quiquampoix, H., & Simonet, P. (2001). Evaluation of biological and physical protection against nuclease degradation of clay-bound plasmid DNA. Applied and Environmental Microbiology, 67(1), 293–299.

Direito, S. O., Marees, A., & Röling, W. F. (2012). Sensitive life detection strategies for low-biomass environments: Optimizing extraction of nucleic acids adsorbing to terrestrial and Mars analogue minerals. FEMS Microbiology Ecology, 81(1), 111–123.

Franchi, M., Bramanti, E., Morassi Bonzi, L., Luigi Orioli, P., Vettori, C., & Gallori, E. (1999). Clay-nucleic acid complexes: Characteristics and implications for the preservation of genetic material in primeval habitats. Origins of Life and Evolution of the Biosphere, 29(3), 297–315.

Harrison, J. B., Sunday, J. M., & Rogers, S. M. (2019). Predicting the fate of eDNA in the environment and implications for studying biodiversity. Proceedings of the Royal Society B: Biological Sciences, 286(1915), 20191409. https://doi.org/10.1098/rspb.2019.1409

Homel, K. M., Franklin, T. W., Carim, K. J., McKelvey, K. S., Dysthe, J. C., & Young, M. K. (2021). Detecting spawning of threatened chum salmon Oncorhynchus keta over a large spatial extent using eDNA sampling: Opportunities and considerations for monitoring recovery. Environmental DNA, 3(3), 631–642.

Jo, T., & Yamanaka, H. (2022). Meta-analyses of environmental DNA downstream transport and deposition in relation to hydrogeography in riverine environments. Freshwater Biology.

Kirtane, A., Atkinson, J. D., & Sassoubre, L. (2020). Design and validation of passive environmental DNA samplers using granular activated Ccarbon and Mmontmorillonite clay. Environmental Science & Technology, 54(19), 11961–11970.

Kirtane, A., Wilder, M. L., & Green, H. C. (2019). Development and validation of rapid environmental DNA (eDNA) detection methods for bog turtle (Glyptemys muhlenbergii). PloS One, 14(11), e0222883.

Klymus, K. E., Merkes, C. M., Allison, M. J., Goldberg, C. S., Helbing, C. C., Hunter, M. E., Jackson, C. A., Lance, R. F., Mangan, A. M., & Monroe, E. M. (2020). Reporting the limits of detection and quantification for environmental DNA assays. Environmental DNA, 2(3), 271–282.

Lamb, P. D., Fonseca, V. G., Maxwell, D. L., & Nnanatu, C. C. (2022). Systematic review and meta-analysis: Water type and temperature affect environmental DNA decay. Molecular Ecology Resources.

Lever, M. A., Torti, A., Eickenbusch, P., Michaud, A. B., Šantl-Temkiv, T., & Jørgensen, B. B. (2015). A modular method for the extraction of DNA and RNA, and the separation of DNA pools from diverse environmental sample types. Frontiers in Microbiology, 6, 476.

Levy-Booth, D. J., Campbell, R. G., Gulden, R. H., Hart, M. M., Powell, J. R., Klironomos, J. N., Pauls, K. P., Swanton, C. J., Trevors, J. T., & Dunfield, K. E. (2007). Cycling of extracellular DNA in the soil environment. Soil Biology and Biochemistry, 39(12), 2977–2991.

Mauvisseau, Q., Harper, L. R., Sander, M., Hanner, R. H., Kleyer, H., & Deiner, K. (2022). The multiple states of environmental DNA and what is known about their persistence in aquatic environments. Environmental Science & Technology, 56(9), 5322–5333.

Nagler, M., Podmirseg, S. M., Ascher-Jenull, J., Sint, D., & Traugott, M. (2022). Why eDNA fractions need consideration in biomonitoring. Molecular Ecology Resources, 22(7). https://doi.org/10.1111/1755-0998.13658

Pawlowski, J., Bruce, K., Panksep, K., Aguirre, F. I., Amalfitano, S., Apothéloz-Perret-Gentil, L., Baussant, T., Bouchez, A., Carugati, L., & Cermakova, K. (2021). Environmental DNA metabarcoding for benthic monitoring: A review of sediment sampling and DNA extraction methods. Science of the Total Environment, 151783.

Pietramellara, G., Franchi, M., Gallori, E., & Nannipieri, P. (2001). Effect of molecular characteristics of DNA on its adsorption and binding on homoionic montmorillonite and kaolinite. Biology and Fertility of Soils, 33(5), 402–409.

Pont, D., Rocle, M., Valentini, A., Civade, R., Jean, P., Maire, A., Roset, N., Schabuss, M., Zornig, H., & Dejean, T. (2018). Environmental DNA reveals quantitative patterns of fish biodiversity in large rivers despite its downstream transportation. Scientific Reports, 8(1), 1–13.

Rees, H. C., Maddison, B. C., Middleditch, D. J., Patmore, J. R., & Gough, K. C. (2014). The detection of aquatic animal species using environmental DNA–a review of eDNA as a survey tool in ecology. Journal of Applied Ecology, 51(5), 1450–1459.

Rodriguez-Ezpeleta, N., Morissette, O., Bean, C. W., Manu, S., Banerjee, P., Lacoursière-Roussel, A., Beng, K. C., Alter, S. E., Roger, F., & Holman, L. E. (2021). Trade-offs between reducing complex terminology and producing accurate interpretations from environmental DNA: Comment on “Environmental DNA: What’s behind the term?” by Pawlowski et al.,(2020). Molecular Ecology.

Saeki, K., Kunito, T., & Sakai, M. (2010). Effects of pH, ionic strength, and solutes on DNA adsorption by andosols. Biology and Fertility of Soils, 46(5), 531–535.

Sakata, M. K., Yamamoto, S., Gotoh, R. O., Miya, M., Yamanaka, H., & Minamoto, T. (2020). Sedimentary eDNA provides different information on timescale and fish species composition compared with aqueous eDNA. Environmental DNA, 2(4), 505–518.

Sassoubre, L. M., Yamahara, K. M., Gardner, L. D., Block, B. A., & Boehm, A. B. (2016). Quantification of environmental DNA (eDNA) shedding and decay rates for three marine fish. Environmental Science & Technology, 50(19), 10456–10464.

Sepulveda, A. J., Schabacker, J., Smith, S., Al-Chokhachy, R., Luikart, G., & Amish, S. J. (2019). Improved detection of rare, endangered and invasive trout in using a new large-volume sampling method for eDNA capture. Environmental DNA, 1(3), 227–237. https://doi.org/10.1002/edn3.23

Sheng, X., Qin, C., Yang, B., Hu, X., Liu, C., Waigi, M. G., Li, X., & Ling, W. (2019). Metal cation saturation on montmorillonites facilitates the adsorption of DNA via cation bridging. Chemosphere, 235, 670–678.

Shogren, A. J., Tank, J. L., Andruszkiewicz, E., Olds, B., Mahon, A. R., Jerde, C. L., & Bolster, D. (2017). Controls on eDNA movement in streams: Transport, Retention, and Resuspension. Scientific Reports, 7(1), 5065. https://doi.org/10.1038/s41598-017-05223-1

Sodnikar, K., Parker, K. M., Stump, S. R., ThomasArrigo, L. K., & Sander, M. (2021). Adsorption of double-stranded ribonucleic acids (dsRNA) to iron (oxyhydr-) oxide surfaces: Comparative analysis of model dsRNA molecules and deoxyribonucleic acids (DNA). Environmental Science: Processes & Impacts, 23(4), 605–620.

Taberlet, P., Coissac, E., Hajibabaei, M., & Rieseberg, L. H. (2012). Environmental DNA. Molecular Ecology, 21(8), 1789–1793. https://doi.org/10.1111/j.1365-294X.2012.05542.x

Torti, A., Lever, M. A., & Jørgensen, B. B. (2015). Origin, dynamics, and implications of extracellular DNA pools in marine sediments. Marine Genomics, 24, 185–196.

Tsuji, S., Takahara, T., Doi, H., Shibata, N., & Yamanaka, H. (2019). The detection of aquatic macroorganisms using environmental DNA analysis—A review of methods for collection, extraction, and detection. Environmental DNA, 1(2), 99–108.

Turner, C. R., Uy, K. L., & Everhart, R. C. (2015). Fish environmental DNA is more concentrated in aquatic sediments than surface water. Biological Conservation, 183, 93–102.

Wickham, H., Averick, M., Bryan, J., Chang, W., McGowan, L. D., François, R., Grolemund, G., Hayes, A., Henry, L., & Hester, J. (2019). Welcome to the Tidyverse. Journal of Open Source Software, 4(43), 1686.

Xu, R., Zhao, A., & Ji, G. (2003). Effect of low-molecular-weight organic anions on surface charge of variable charge soils. Journal of Colloid and Interface Science, 264(2), 322–326.

Ye, J., Coulouris, G., Zaretskaya, I., Cutcutache, I., Rozen, S., & Madden, T. L. (2012). Primer-BLAST: A tool to design target-specific primers for polymerase chain reaction. BMC Bioinformatics, 13, 134. https://doi.org/10.1186/1471-2105-13-134

Yu, W. H., Li, N., Tong, D. S., Zhou, C. H., Lin, C. X. (Cynthia), & Xu, C. Y. (2013). Adsorption of proteins and nucleic acids on clay minerals and their interactions: A review. Applied Clay Science, 80–81, 443–452. https://doi.org/10.1016/j.clay.2013.06.003

Yuan, Q.-B., Huang, Y.-M., Wu, W.-B., Zuo, P., Hu, N., Zhou, Y.-Z., & Alvarez, P. J. (2019). Redistribution of intracellular and extracellular free & adsorbed antibiotic resistance genes through a wastewater treatment plant by an enhanced extracellular DNA extraction method with magnetic beads. Environment International, 131, 104986.

