## Supporting information for "Can we sort states of environmental DNA (eDNA) from a single sample?"

Table S1: NCBI Accession numbers of sequences used to design the qPCR assays in this study

| **Target species** | **Target gene** | **Accession numbers** |
|---|---|---|
| Clum salmon (Oncorhynchus keta) | Mitochondrial Cytochrome Oxidase subunit 1 (COX1) | MN850432.1, MT577143.1-MT577155.1, MK216596.1, MK991792.1, KX145589.1, KX145300.1, KX145283.1, KX145253.1, KX144995.1, KX958413.1, KU756203.1-KU756205.1, LC094464.1 - LC094479.1, KR778858.1, KR778851.1, KR778852.1, MZ098006.1, MZ098005.1, ON000117.1, MN756459.2, MN756291.2, OL674493.1 |
| Chicken (Gallus gallus) | Mitochondrial NADH Dehydrogenase 2 (ND2) | MT773644.1 |
| Mouse (Mus musculus) | Mitochondrial NADH Dehydrogenase 2 (ND2) | MT937073.1 |
| Mouse (Mus musculus) | Nuclear Transforming Growth Factor beta 1 (TGFb1) | NC000073.7 |

Table S2: gBlock sequences used this study. Position of Primers and Probes are marked as follows: Forward Primer, Probe, Reverse Primer (reverse compliment).

| Target species | Target gene | gBlock sequence |
|---|---|---|
| Mouse (*Mus musculus*) | ND2 | GTAAGGTCAGCTAATTAAGCTATCGGGCCCATACCCCGAAAACGTTGGTTTAAA<br>TCCTTCCCGTACTAATAAATCCTATCACCCTTGCCATCATCTACTTCACAATCTTC<br>TTAGGTCCTGTAATCACAATATCCAGCACCAACCTAATACTAATATGAGTAGGC<br>CTGGAATTCAGCCTACTAGCAATTATCCCCATACTAATCAACAAAAAAAACCCA<br>CGATCAACTGAAGCAGCAACAAAATACTTCGT |
| Mouse (*Mus musculus*) | TGFb1 | GGCGGTGGTGGTGCACGTCTTCAATTCCAGCACTCAGGAGGCAGAGGCAGGTGG<br>ATCTCTATGAGTGCGAGTCCAACCAGGCTTAAAACGTCCTGCCTGGACTAGGCT<br>GGCTTCAAACTTGCAGCGATCCTCCCATTTCGGCTTCTTGACTACAGTTGTGCAC<br>AGAGGTGCATGCCTGCCTGGTTCACATGTGTCAAGTTTTATTTTTTATTATTTTCT<br>TTTACATGCATGTTTTGTCTTTGTGGATGTA |
| Chicken (*Gallus gallus*) | ND2 | ACCATTGAATCTTAGCCTGAACAGGCTTAGAGATCAACACCTTAGCCATCATCCC<br>CCTCATCTCCAAGTCACACCACCCCCGAGCGATTGAAGCCACTATCAAATATTTC<br>CTCACCCAATCAACTGCATCAGCCCTAATCCTCTTCTCGAGCATAACCAACGCCT<br>GATCCACCGGACAATGAGACATTACACAACTAAACCACCCGACATCATGCCTAA<br>TATTAACAATAGCAATCGCAATCAAATTAGG |
| Chum salmon (*Oncorhyncus keta*) | COI | TTATTACGACCATTATCAACATAAAACCCCCAGCTATTTCTCAGTACCAAACCCC<br>GCTTTTTGTCTGAGCTGTACTAATCACTGCTGTACTTCTACTATTATCACTCCCCG<br>TTCTGGCAGCAGGTATTACTATGTTGCTCACAGATCGAAATTTAAACACCACTTT<br>CTTTGACCCAGCGGGCGGCGGAGATCCAATTTTATACCAACACCTCT |

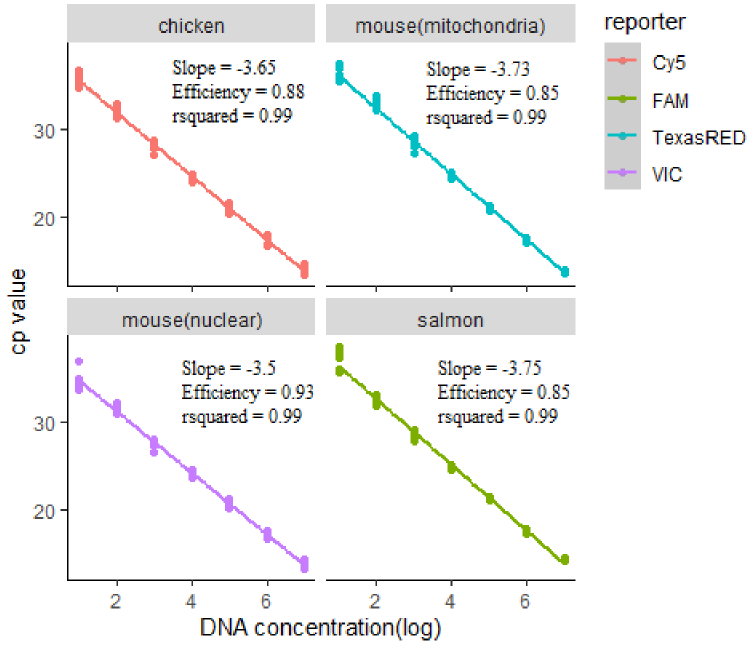


Figure S1: Pooled standard curves from all qPCR plates used to quantify the DNA concentration of the four targets in the experiment.

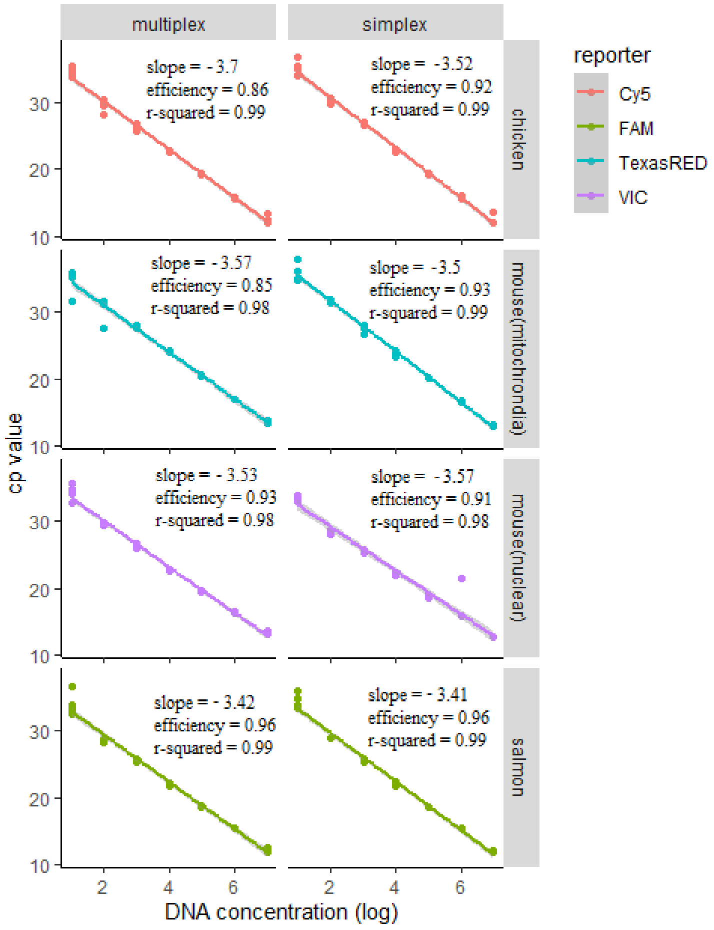


Figure S2: Evaluation of qPCR standard curves performed in both simplex and multiplex reaction types using gBlocks from four targets used in this experiment. Each target is represented by a unique fluorophore.

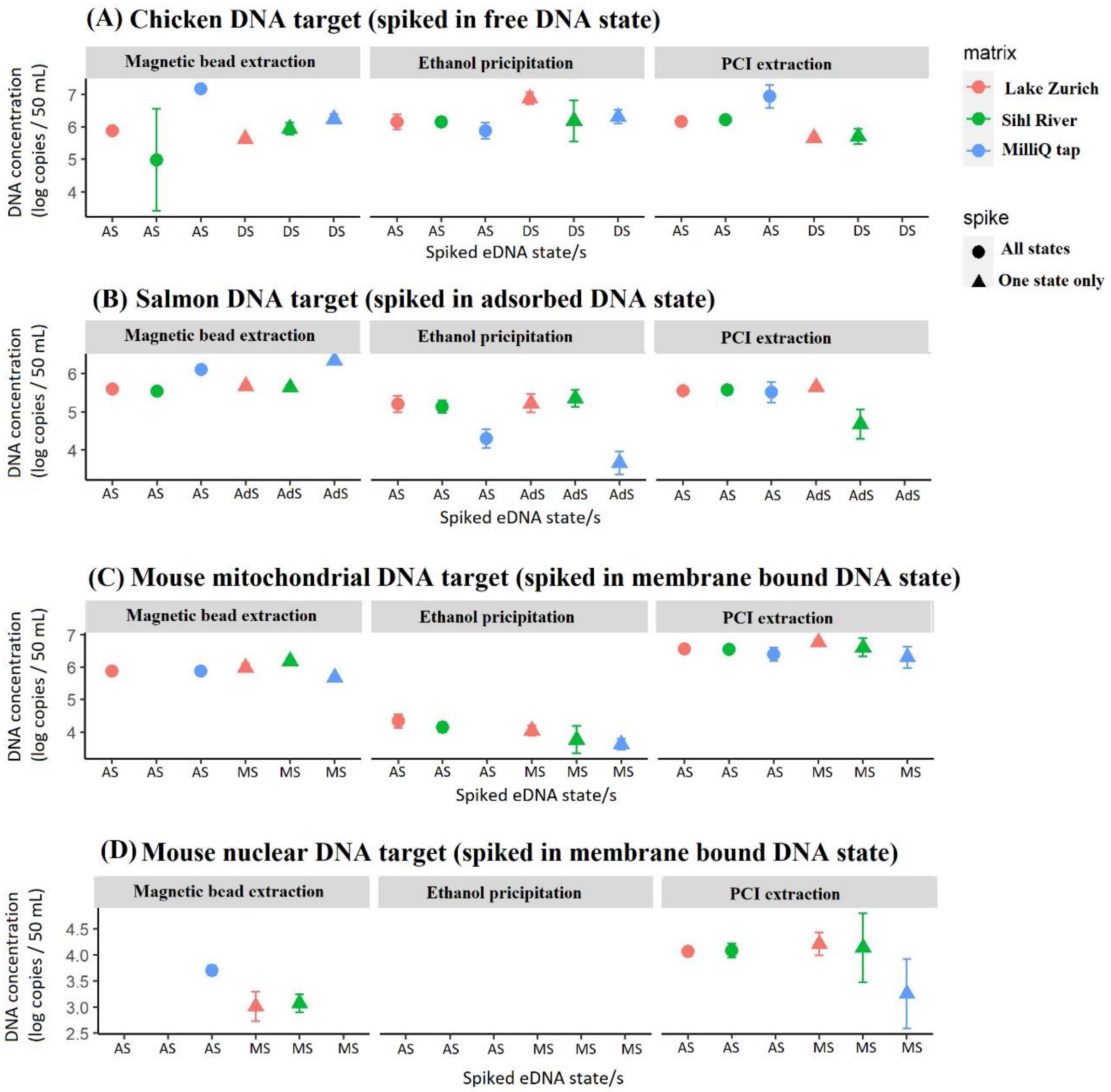


Figure S3: DNA concentration (log copies / 50 mL) of spiked eDNA states observed in three different extraction methods. Colors represent the water matrix type while the shape of the data points indicates whether all states were spiked together or the target state spiked by itself. The x-axis shows the spiked eDNA state (AS = All States, DS = Dissolved State, AdS = Adsorbed State, and MS = Membrane-bound State). Error bars indicate the standard deviation between three biological replicates.

Table S3: Breakdown of spike, recovery, and loss of target DNA based on the state the DNA was spiked in and extraction method.

| | Dissolved State (copies/50ml) | Mitochondrial membrane-bound state (copies/50ml) | Nuclear membrane-bound state (copies/50ml) | Adsorbed state (copies/50ml) |
|---|---|---|---|---|
| Spike concentration | $3.9*10^{8}$ | $8.5*10^{6}$ | $3.0*10^{4}$ | $2.5*10^{7}$ |
| Ethanol Precipitation | $2.9*10^{6} \pm 2.7*10^{6}$ | $1.6*10^{4} \pm 7.1*10^{3}$ | BLOQ | $1.3*10^{5} \pm 9.8*10^{4}$ |
| Magnetic bead extraction | $3.3*10^{6} \pm 5.7*10^{6}$ | $9.2*10^{5} \pm 4.1*10^{5\#}$ | $2.6*10^{3} \pm 2.3*10^{3\#}$ | $8.6*10^{5} \pm 7.5*10^{5}$ |
| Phenol-chloroform extraction | $3.2*10^{6} \pm 4.8*10^{6}$ | $3.8*10^{6} \pm 1.3*10^{6}$ | $1.6*10^{4} \pm 7.9*10^{3}$ | $3.3*10^{6} \pm 1.5*10^{5}$ |
| Total recovered | $8.8*10^{6} \pm 9.3*10^{6}$ | $4.6*10^{6} \pm 1.6*10^{6}$ | $1.4*10^{4} \pm 8.6*10^{3}$ | $1.3*10^{6} \pm 5.7*10^{5}$ |

BLOQ refers to Below Limit of Quantification

### Some replicates of this group were Below Limit of Quantification
